# M13 phage display to identify a permeating peptide against hyperconcentrated mucin

**DOI:** 10.1101/659573

**Authors:** Jasmim Leal, Tony Dong, Feng Gao, Melissa Soto, Hugh D.C. Smyth, Debadyuti Ghosh

**Author notes:** Division of Molecular Pharmaceutics and Drug Delivery, College of Pharmacy, The University of Texas at Austin, 2409 University Ave., STOP A1920, Austin, TX 78712, USA.

## Abstract

Mucus is an impregnable barrier for drug delivery across the epithelia for treatment of mucosal-associated diseases. While current carriers are promising for mucus penetration, their surface chemistries do not possess chemical complexity to probe and identify optimal physicochemical properties desired for mucus penetration. As initial study, we use M13 phage display presenting random peptides to select peptides that can facilitate permeation through hyperconcentrated mucin. Here, a net-neutral charge, hydrophilic peptide was identified to facilitate transport of phage and fluorophore conjugates through mucin barrier compared to controls. This initial finding warrants further study to understand how composition and spatial distribution of physicochemical properties of peptides can be optimized to improve transport across the mucus barrier.

## INTRODUCTION

Mucosal surfaces of gastrointestinal, vaginal, respiratory and nasal tissues provide innate protection from pathogens and allow select passage of nutrients during homeostasis (1–3). However, in drug delivery across these mucosal-associated tissues and in diseases of the mucus environment such as cystic fibrosis or chronic obstructive pulmonary disease, mucus serves as a physical barrier to prevent drug transport for successful therapy and also creates a local environment to trap and protect pathogens resulting in chronic bacterial infections (1). Mucus, comprising of mainly mucin glycoprotein (> 95% mucin in all mucus types), is a viscous gel that binds drugs via intermolecular interactions whereby preventing effective penetration. Mucin consists of repeated proline-threonine-serine domains with hydrophilic, glycosylated areas interspaced with hydrophobic globular regions rich in cysteines. As a result, drugs including positively charged tobramycin and gentamicin bind to the net-negatively charged oligosaccharides on mucins, and lipophilic drugs such as progesterone and testosterone interact with hydrophobic domains via van der Waals interactions (4–7). Penetration through the mucus barrier would enable effective delivery of drugs towards successful treatment of mucosal-associated diseases. Technologies including permeation enhancers (8), transporter conjugates (9), mucoadhesive polymers (10), and micro- and nanoparticle systems (11) have been developed to facilitate transport across the mucus, yet it remains a challenge to achieve transmucosal delivery at therapeutic doses. Recent work has shown that hydrophilic, net-neutral formulations improve transport through mucus barriers (12–21). In particular, particles densely coated with low molecular weight poly(ethylene) glycol (PEG) have demonstrated rapid transport through mucus barriers including cervicovaginal mucus and mucus in pulmonary and gastrointestinal tissues (12–18, 20, 21). Hydrophilic PEG circumvents binding to the hydrophobic domains on mucus and these formulations possess neutral charge, which avoids electrostatic interactions with negatively charged mucin (20). While promising, PEG-functionalized formulations have potential challenges including unexpected immunogenicity and hindered cellular internalization and escape, which highlight the need for additional alternatives. Despite their long systemic circulation half-life, PEG formulations have been found to be rapidly cleared after repeated dosing due to the production of anti-PEG antibodies, thereby limiting its use for repeated dosing regimens (22–25). PEGylation also has been shown to impair cell internalization and intracellular drug release (26–28). DNA polyplexes coated with PEG demonstrated 10-fold reduction in uptake in cell lines compared to non-functionalized polyplexes, and 100 to 1000-fold decrease in release of reporter luciferase gene relative to equivalent naked DNA delivery (26). PEG is thought to impair the “proton sponge effect” whereby the influx of hydrogen ions into the endosomes and lysosomes causes organelle rupture and allows for drug escape (26, 27). These challenges of PEGylated formulations present substantial bottlenecks to their long-term success for transmucosal delivery. In addition, formulations functionalized with a single PEG coating will possess uniform surface chemistries, which dictate and limit the number of potential interactions (or lack thereof) to query the complex mucus environment. While mucus is net-negative charge and is hydrophilic, there are spatially heterogeneous regions of positive and negative charges and hydrophilic and hydrophobic domains. Mucus acts an “interaction filter” able to form interactions with different substrates (16, 29, 30). From these observations, it is critical to expand the repertoire of carriers or substrates with greater chemical complexity and diversity of physicochemical properties that achieve unhindered transport through the mucus barrier.

Biomolecules such as viruses, proteins, and peptides possess the genetic, chemical, and physical properties for successful transport and transmission through mucosal epithelia. Protein coated viruses (i.e. capsid viruses) have been shown to diffuse through gastric and nasal mucus barriers to effectively bind and infect underlying epithelia. Norwalk virus-like particles (38 nm diameter) and human papilloma virus-like particles (55 nm diameter) transport through healthy mucus as rapidly as in saline (31). They demonstrated significantly enhanced transport compared with similar-sized (59 nm) synthetic polystyrene nanoparticles, in part due to their complex, dense coating of net-neutral, alternating positive and negatively charged amino acid residues (1, 32). In addition, recent work indicates that molecules with asymmetric charge properties improve transport through hydrogels and mucus, which underscores the importance of the distribution of physicochemical properties on mucus interactions. Li et al. demonstrated that small-sized peptides spatially arranged with a block of positive and then negative charged amino acids (AKAKAKAKAKAEAEAEAEAE) improved transport through mucus than in control saline and diffused better than only positive or all negatively charged peptides (33). Further, block peptides with same net charge as alternately charged peptides (AKAEAKAEAKAEAKAEAKAE) demonstrated differences in diffusive transport through mucin. Block peptides had different diffusivities in saline and mucin, whereas alternately charged peptides with the same overall charge demonstrated negligible change in transport with or without the presence of mucin. The mucin barrier is able to delineate the local differences in charge distribution from amino acids on the nanometer scale (33). This finding emphasizes how spatial distribution of amino acids (and their functional groups) can greatly impact transport behavior through the mucus barrier. As a result, substrates with combinations of physicochemical properties can be used to screen and identify optimized, desired functionalities for mucus penetrating delivery.

Towards that end, the goal of this work is to use bacteriophage libraries as complex substrates to screen and identify peptides that facilitate transport through mucus barriers. Bacteriophage (phage), or viruses that infect bacteria, have been developed as display systems able to express recombinant peptides or proteins on their viral coat proteins (i.e. capsid proteins). One particular phage, filamentous M13 (Figure 1; ~6 nm diameter, 880 nm length) is genetically modifiable such that peptides can be displayed on the various coat proteins of the virus (e.g. p3 and p8 as shown in Figure 1). By genetically engineering random DNA oligonucleotides encoding peptides into the phage genome, it is feasible to generate phage-presenting random peptide libraries (on a single coat protein) with diversity of up to 10^9^ – 10^10^ different peptides (i.e. different peptide per phage). The phage libraries have peptides with diverse functionalities due to the side chain groups of their amino acids. The libraries are incubated against specific targets such as cells or biomolecules (34), and by an iterative process of collection, amplification and re-incubation, peptides with affinity for the target are selectively enriched and identified from the library.

**Figure 1.**
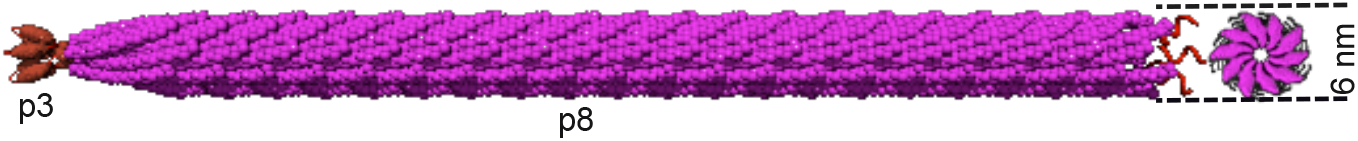
Cartoon of filamentous M13 bacteriophage with minor coat protein p3 at the proximal end and p8 major coat protein along the length of the bacteriophage.

Using these phage libraries and applying an iterative screening strategy against mucus, peptides can be selected that are muco-inert and mucus-penetrating without *a priori* knowledge of the heterogeneous mucus composition. Additionally, this approach allows us to screen peptides that transport through mucus irrespective of the heterogeneous microscale rheology of the mucin (35). The large diversity of random phage-presenting peptides is essentially a collection of formulations with permutations of physicochemical properties. Here, we have identified select peptides that exhibit improved diffusivity in hyperconcentrated mucin compared to controls, and from phage screening, have identified physicochemical properties that are critical for transport across the barrier. This study will serve as the initial step towards elucidating the role of composition towards mucus penetration and importantly, help advance the development of new mucus-penetrating drug delivery systems.

## MATERIALS AND METHODS

### Phage Libraries

The Ph.D. 7-mer random peptide phage library (New England Biolabs (NEB), Ipswich, MA) was used to identify peptides that facilitate transport through mucin. The Ph.D. library displays random peptides 7 amino acids in length on the N-terminus of p3 coat protein of M13 bacteriophage. M13 displays peptides encoded by [NNK]_7_ with a flexible GGGS linker between Acc65I and EagI restriction endonuclease sites, where N = A, C, G, or T and K is G or T to remove two out of three potential stop codons while maximizing sequence diversity. The Ph.D. library was calculated to have diversity of 2*10^9^ (i.e. different peptides displayed on each phage), as provided from manufacturer. To visualize phage on agar plates, the M13 vector also contains *lacZα* gene, where upon bacterial infection with M13 phage, the resulting plaques will appear blue when plated on agar with X-gal substrate.

### Phage Screening Assay

To use phage libraries in mucin and identify clones able to diffuse through mucin, we developed a phage screening assay (Figure 2). 10^11^ plaque forming units (pfu)/mL Ph.D. 7-mer phage library were incubated with reconstituted mucin (see **Mucin Preparation** in **Materials and Methods**) in the apical chamber of a 12 mm diameter (12 well) polyester Transwell chamber with 3 micrometer pore size (Corning, Corning, NY). Before addition of phage and mucin to the apical chamber, the basolateral chamber (i.e. donor compartment) was filled with phosphate buffered saline without calcium and magnesium (PBS, Corning, Corning, NY), as recommended by manufacturer. After 1 hour, peptide-presenting phage that diffused and penetrated through mucin were collected in the donor compartment. To make more copies of collected phage, they were incubated with early log phase XL-1 Blue *E. coli* (OD_600_ = 0.2-0.4; Agilent, Santa Clara, CA) in 20 mL Luria-Bertani culture media (LB) for 30 minutes non-shaking to allow for infection and then grown for 3 hours at 37°C and 225 rpm for amplification (MaxQ 4000, ThermoFisher Scientific, Hannover Park, IL). After, the culture was centrifuged at 10,000 rpm for 15 min (Sorvall Legend XFR, Newtown, CT) to separate XL-1 *e. coli* from phage in the supernatant. Following standard techniques, phage were then precipitated overnight at 4°C with 1/6 volume poly(ethyelene) glycol 8000 with 2.5 M sodium chloride (PEG 8000/2.5 M NaCl) (Fisher BioReagents, Pittsburgh, PA) (36). Samples were centrifuged at 10,000 rpm for 15 min and the resulting phage pellet was resuspended in 1 mL PBS. To remove residual bacteria and debris, samples were recentrifuged at 13,000 rpm for 2 minutes using a ThermoScientific Legend Microcentrifuge (Hannover Park, IL). Phage supernatant were transferred to a fresh microcentrifuge tube and as a final purification step, samples were then re-precipitated with 1/6 volume PEG 8000/2.5 M NaCl and stored on ice for 1 h. Samples were then centrifuged at 10,000 rpm for 15 minutes, and the purified phage pellet was resuspended in 200 μL PBS. Adapting from the protocol by NEB, phage were quantified by titration, i.e. phage activity counted by formation of plaques. Briefly, serial ten-fold dilutions of amplified phage were prepared, and 10 μL from serial dilutions were incubated each with 200 μL mid-log XL-1 (OD_600_ = 0.5-0.8) for 1-5 minutes to allow sufficient time for infection, mixed with 1 mL warm liquid agar and overlaid on 6-well solid agar plates. Plaques indicative of phage-infected bacteria were counted by standard *lacZα* blue screening and quantified as plaque forming units (pfu). After titering amplified phage from initial round of screening, three additional rounds of mucin-penetration screening were subsequently performed using equivalent amounts of phage per round. After four rounds of screening, mucin-penetrating phage were titered without amplification on 6-well LB-agar plates, and individual clones were isolated for DNA sequencing and peptide identification. Individual plaques were isolated and grown in 5 mL of early-log XL-1 Blue (OD_600_ = 0.2-0.4) overnight (~16 hours) at 37°C and 225 rpm, and phage DNA from infected bacteria was isolated using Qiagen QiaPrep Spin Mini Kit (Qiagen, CA) following manufacturer’s protocol. Sanger sequencing of isolated clones was done at the DNA sequencing facility at the Institute for Cellular and Molecular Biology at The University of Texas at Austin.

**Figure 2.**
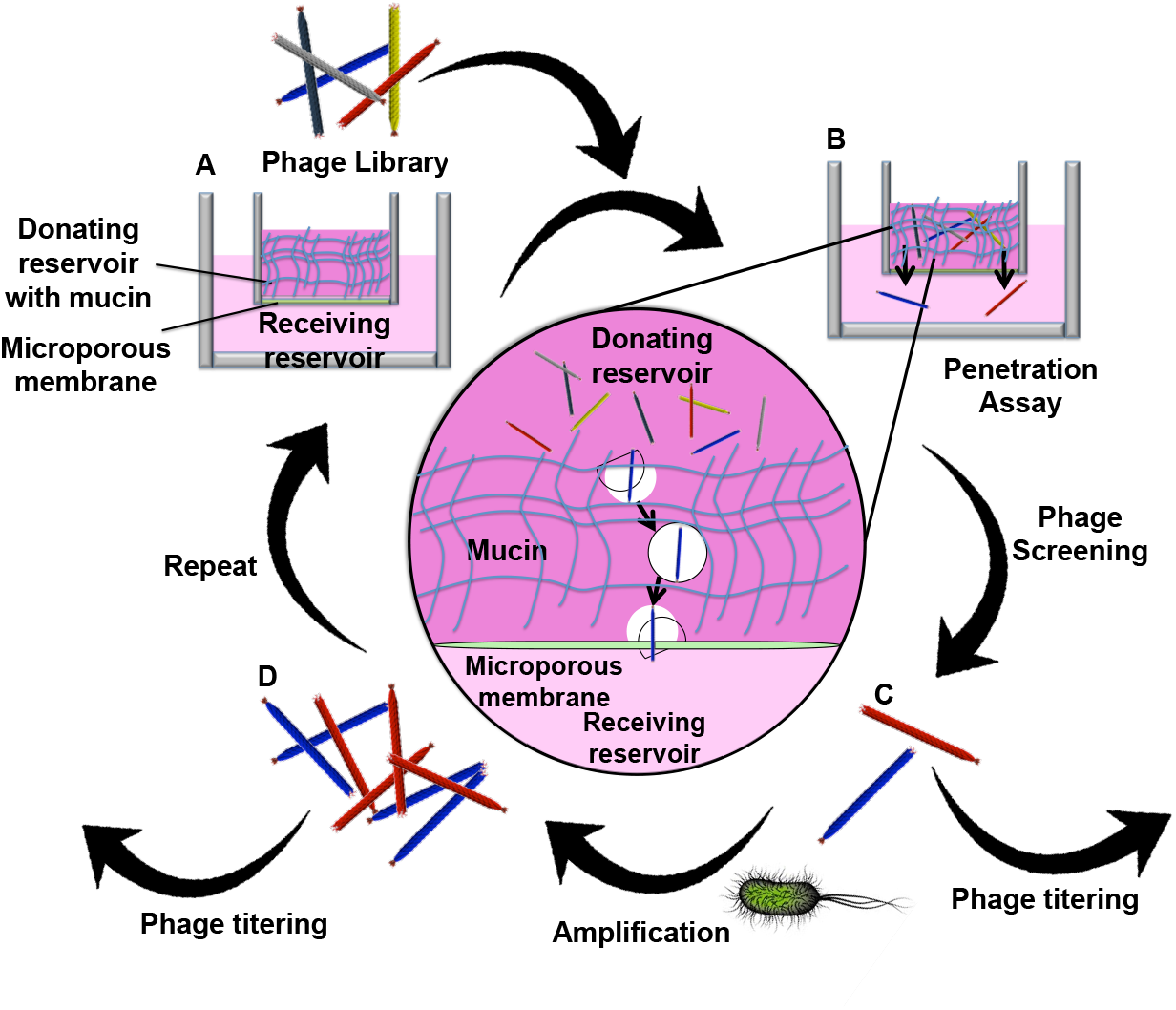
Schematic of phage screening assay. **A.** linear Ph.D. M13 library is incubated with hyperconcentrated mucin in the donating reservoir of receiving chamber. **B.** during timecourse, select phage diffuse through the mucin barrier into the receiving reservoir (see zoom in center). **C.** select phage are collected and are split-quantified by titering (i.e. double layer plaque assay) or amplified in E. coli to generate more phage that eluted through mucin. **D.** Amplified phage are quantified by titering and then the iterative selection, or biopanning, is repeated in the next round. This assay is done in multiple rounds (at least three).

### Mucin Preparation

Type II mucin (Sigma Aldrich, Saint Louis, MO) was used for reconstitution of mucin. As a simple model, 8% mucin preparation (w/w) was used to recapitulate high concentrations observed in cystic fibrosis patients (37, 38). Lyophilized mucin was reconstituted in PBS at pH= 7.2 following a protocol from McGill and Smyth (39).

### In silico analysis of physicochemical properties

The net charge at pH = 7 of sequences was calculated using Protein Calculator v3.4 (Scripps Research Institute, La Jolla, CA). GRAVY (grand average of hydropathy) was calculated according to Kyte-Doolittle (40) using www.gravy-calculator.de.

### Validation Transport Assays

To validate clones could facilitate improved transport, transport was compared between selected clones with negative control M13 that does not display any recombinant peptide on p3 (designated as “wild-type”). Adapting from the transwell assay described earlier, 1*10^10^ pfu of each phage was incubated in separate transwells with 8% mucin and incubated for 1 h in the apical chamber. After, the donor compartment was collected (1.5 mL PBS) and quantified by titering to count mucin-penetrated phage. Samples were run in triplicate and in multiple experiments. Error was calculated as standard deviation (s.d.).

### Bulk Diffusion Assays and Calculation of Diffusivities

To quantify effective diffusivities of phage, a transwell assay adapted from above was performed. Equivalent amounts of clones and control M13 phage were incubated in 8% mucin in the apical chamber of multiple transwells for 15, 30, 45, and 60 minutes (each transwell is a specific timepoint). At each timepoint, penetrating phage were collected from the basolateral chamber and titered. Samples were run in triplicate. To calculate diffusivity of phage through mucin, a method developed by Hu et al. for phage diffusion through gels and biofilms (41) and confirmed by others (42) was used. Fick’s first law of diffusion was used to determine the apparent diffusion coefficient (D_app_). Assuming boundary conditions surrounding the membrane are not involved,

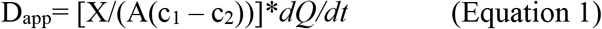

where Q is the amount of solute transferred, c_2_ is concentration in the receiver chamber (basolateral), c_1_ is concentration in the donor chamber (apical), X is the thickness of the membrane, and A is the surface area of the membrane. The results presented is the average of three independent experiments. Hu et al. confirmed the diffusion method for small molecules including D-glucose and both lytic and filamentous phage similar to M13 in mimics of mucus gels and biofilms (41, 43).

### Statistical Analysis

Statistical significance was considered when p < 0.05. Data are presented with standard deviation (s.d.).

## RESULTS

### Enrichment of Phage with Screening Through Mucin

The goal was to establish an assay that allows for screening of phage-based formulations and identify peptides that allow phage transport through a model of reconstituted mucin hydrogel. By setting up the assay that allowed for multiple rounds of screening through mucin, we can reduce the number of phage peptides that penetrate through the mucin barrier. The hyperconcentrated mucin effectively serves as a barrier for phage that penetrate. After the initial round of screening, phage that penetrate through are collected and amplified. By repeating this process for several rounds, phage that nonspecifically diffused due to the heterogeneous pore size of mucin hydrogels can be removed in the additional rounds of screening. Additionally, in each round of screening it is expected that reconstituted mucin would have different microscale viscosity as suggested/demonstrated by others (35); as a result, by subjecting the libraries to repeated screening, phage that penetrate mucin irrespective of the heterogeneity of the composition or pore size would be discovered. This strategy is relevant to human pathology, where there is patient to patient variability in mucus concentration and composition, as often seen in patients with cystic fibrosis or chronic obstructive pulmonary disease (1). For each round of screening, or panning, we incubated 1*10^11^ plaque forming units (i.e. number of infective phage) in freshly prepared 8% mucin and collected the eluates that penetrated after 1 h. We amplified phage eluates to make more copies of the resulting sub-library and iterated this experiment for three additional rounds. To validate enrichment, equivalent amounts of collected phage that were amplified from each round were incubated with mucin and penetrating phage were collected and titered. There is an approximate three-fold improvement, or enrichment in collected phage from phage pool in round three compared with earlier rounds (Figure 3).

**Figure 3.**
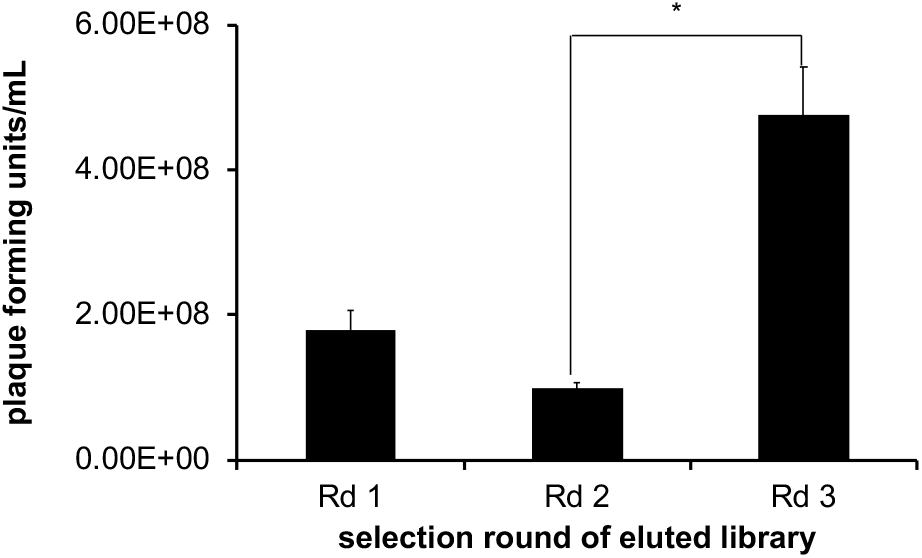
Phage library output (in plaque forming units/mL) after rounds of selection in 8% mucin. At third round, there is enrichment of phage penetrating through mucin (*p < 0.05).

### Identification of Sequences and Their Physicochemical Properties

To identify potential penetrating phage clones, we isolated individual phage after three rounds of screening against mucin. After the third round of screening, penetrating phage were plated on agar, and individual plaques representing individual clones were grown with *E. coli* in culture, and the DNA of the individual clones were isolated. Since the DNA sequence of the M13 vector links with the phenotype, the DNA sequence fused with gene 3 encodes for the peptide displayed on N-terminus of p3 coat protein. After DNA sequencing of 13 clones, we found that two sequences, showed up repeatedly, with sequence TVRTSAD having a frequency of 4/13 clones and NTGSPYE having a frequency of 1/13 (**Table 1**). These sequences have an approximate net charge of 0 (−0.1) and −1 (−1.1), respectively (**Table 1**). In addition, these peptides are hydrophilic, as indicated by the negative GRAVY scores calculated from the Kyte-Doolittle hydropathy plot (**Table 1**)(40, 44). The negative values, which represent the average score of the amino acid at that position relative to surrounding amino acids (i.e. window range), reflects the overall hydrophilic property of the sequence. Also from sequencing, there were clones without insert (wild-type) and sequences with deletions and frame shifts in the randomized 7mer sequence. Since TVRTSAD sequence was most abundant, subsequent experiments were done using this clone.

### Enhanced Transport and Diffusivity of Selected Phage and Peptide Clones

To confirm that identified peptides present on M13 have minimal mucin binding to permit mucin penetration, we tested their transport through mucin compared with negative control wild-type M13. Similar to the screening assay, we incubated equivalent amounts of selected clones and wild type in separate transwells with mucin, collected fractions in the donor compartment, and quantified the fractions by titering. As seen in Figure 4A after 1 h incubation, significantly more TVRTSAD-presenting phage transports through mucin compared with wild-type, suggesting that the peptide improves transport.

**Figure 4.**
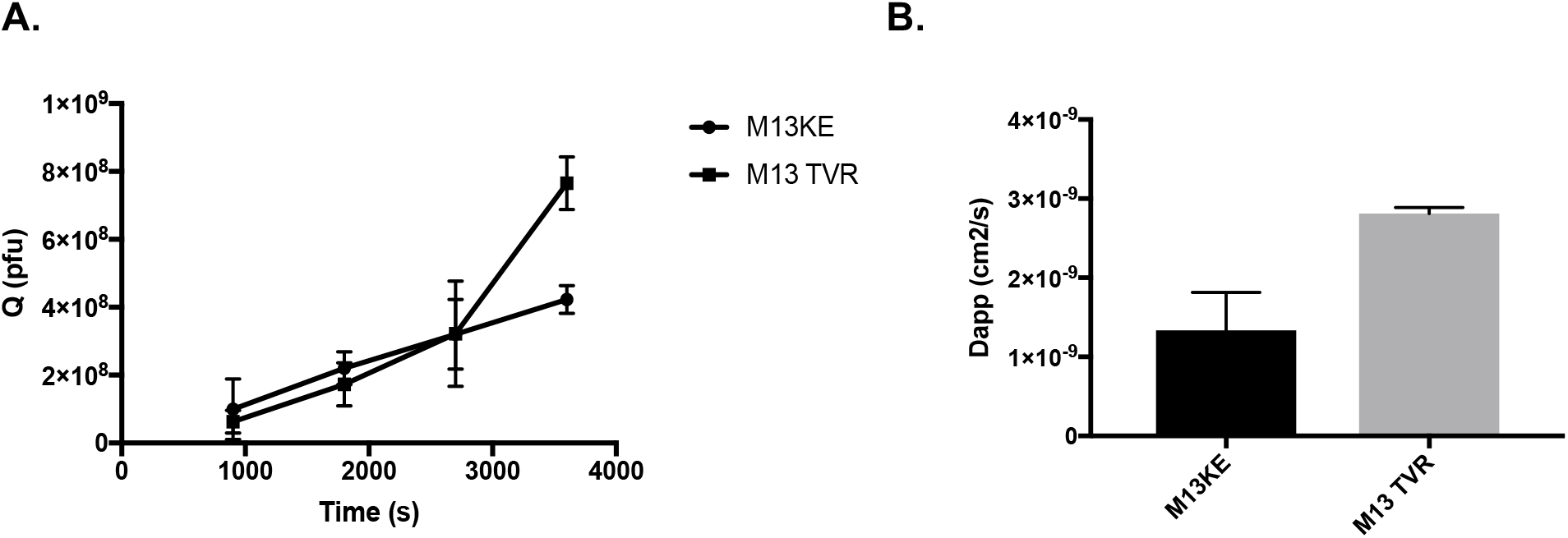
**A.** Amount Q of negative control M13 and TVRTSAD peptide-presenting phage diffusing through mucin at given timepoints. **B.** Apparent diffusivity of negative control M13 and TVRTSAD phage calculated from values taken from **A** and Equation 1.

After validating improved transport with selected phage, we quantified the diffusive behavior of phage in mucin. Previously, the diffusive behavior of lytic and lysogenic filamentous bacteriophage had been quantified through gels and artificial biofilms to determine their feasibility as natural antimicrobial therapeutics to penetrate and effectively diffuse and bind to these complex ecosystems (43). Adapting that method, we quantified diffusivity of our clones compared to controls. In four separate transwells designated for 15, 30, 45, and 60 min of diffusion, either TVRTSAD or wild-type control at equal starting concentrations were incubated for the given timepoint. After each timepoint, samples were collected in the donor compartment and the concentrations were determined by titering. From the amount Q (in plaque forming units), we calculated the apparent diffusivity, or D_app_, of the phage. TVRTSAD-phage has an apparent diffusivity of 2.81*10^−9^ cm^2^/s, which is approximately over two-fold greater than wild-type control with 1.33*10^−9^ cm^2^/s (Figure 4).

Next, we wanted to take the peptides out of the structural context of phage and confirm that the selected peptides facilitate diffusion of molecular conjugates in mucin. Synthetized peptides conjugated with fluorescein isothiocyanate (FITC) were incubated in PBS and mucus and their bulk diffusion was measured; from these experiments, their diffusion coefficients were calculated. The graphs representing amount of solute Q transferred *versus* time in saline and mucus are shown in Figure 5A-C (here, solutes are FITC-peptides and the control fluorescein sodium salt). The amount of solute transferred increased with time. As indicated from the graphs for all solutes, there is a linear relationship between Q and time, and therefore application of Equation 1 is valid for the duration of the experiment. From these measurements, we can calculate their diffusion coefficients. Here, FITC-labeled TVRTSAD had an approximate diffusivity of 5.10*10^−8^ and 0.94*10^−8^ cm^2^/s in PBS and mucin, respectively (Figure 5). While the values are comparable to fluorescein sodium salt in PBS, there is two-fold less diffusion in mucin. Interestingly, the selected TVRTSAD peptide demonstrated improved diffusivity than negative control hepta-arginine (R7; +7 net charge) in mucin (Figure 5D; **Table 2**). Table 2 lists the calculated diffusivities of FITC-peptides and fluorescein salt in mucus and saline. Mucin hinders the diffusion of FITC-peptides and control fluorescein salt. Diffusion coefficients in mucus were decreased almost six-fold compared to PBS (Table 2), which are consistent with prior work where solutes with various molecular weights and physicochemical properties presented similar range of decreased diffusivities in porcine gastric mucus (45). Peptide conjugates demonstrate less effective diffusivity (D_PBS_/D_mucin_) than fluorescein salt but greater effective diffusivity than control peptide R7. TVRTSAD had less than two-fold decrease in diffusion than fluorescein sodium salt in mucin, in spite of being four times large (in molecular weight; **Table 2**).

**Figure 5.**
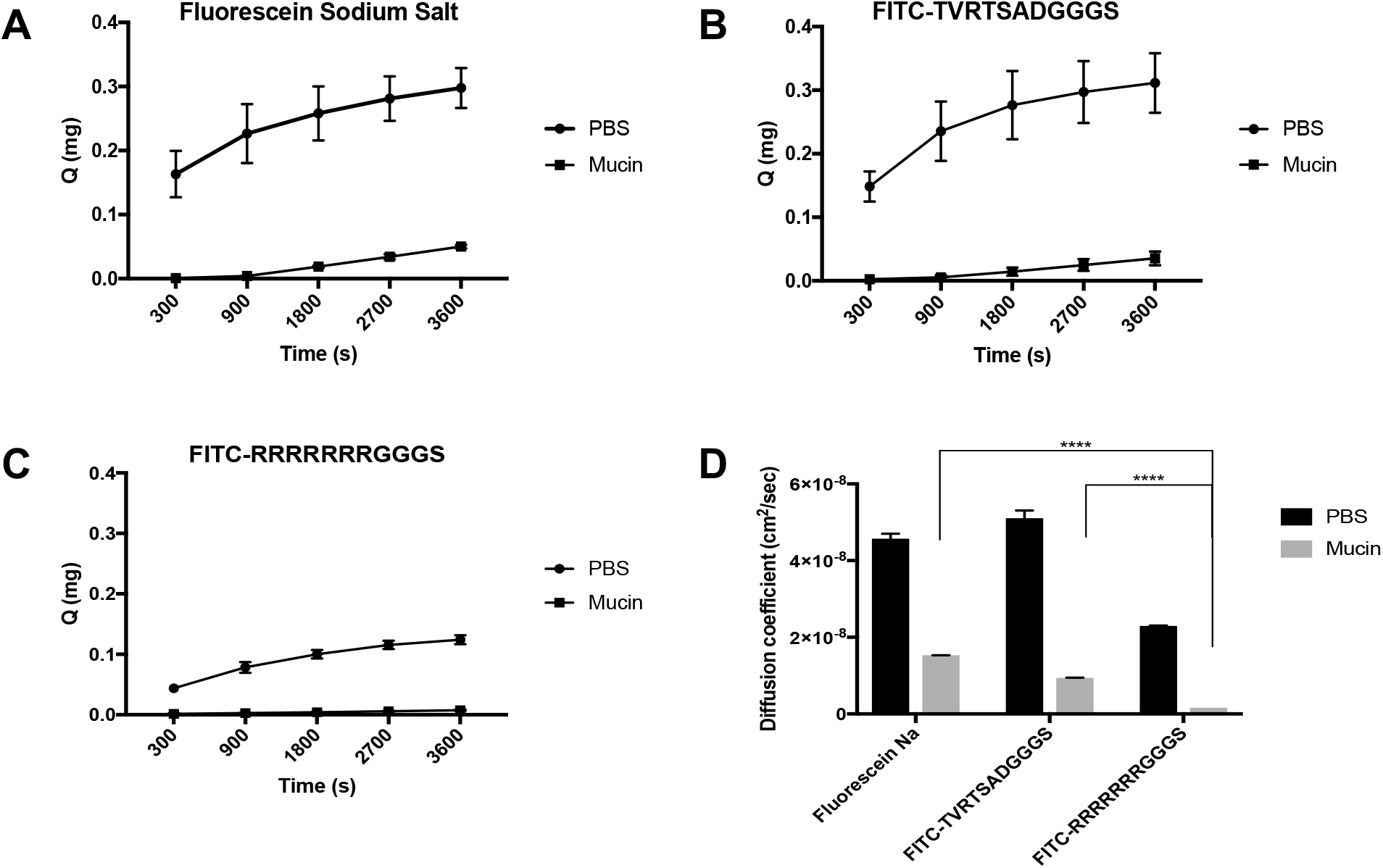
**A-C.** Amount of solute (fluorescein salt, FITC-conjugated TVRTSAD peptide and control R7 peptide) diffusing through mucin and saline (PBS) at given timepoints up to 1 hour. **D.** Calculation of apparent diffusivity of small molecule fluorescein and its peptide conjugates from values taken from **A-C** and Equation 1.

## DISCUSSION

For each successive round of panning, there is an increase in the amount of phage diffusing through the mucin barrier, which suggests that there is enrichment for mucin-penetrating phage (Figure 3). As expected, there is no significant difference of phage going through from pools collected from rounds 1 and 2. This finding is expected, as in the first two rounds in panning against other molecules, including proteins, cells or tissues, conventional panning results in minimal enrichment until at least round three (46). Enrichment is most likely due to the reasoning that after several rounds, non-specific clones were trapped by the mucin gel and with multiple rounds of amplification, more copies of penetrating phage clones were produced and applied in subsequent rounds of screening. While additional rounds of panning could result in further enrichment, it is important to minimize the presence of false positives or parasitic sequences, i.e. phage that amplify rapidly and do not contribute to improved transport, such as wild-type phage (47). Even after three rounds, there is the presence of wild-type, insertless phage, along with other mutant sequences (**Table 1**). Consequently, after three rounds of screening, we proceeded to identify and validate individual clones for enhanced transport.

From *in silico* analysis, we identified the physicochemical properties of select peptides from library panning. The most abundant peptide sequence, TVRTSAD, was neutral and hydrophilic. This result may be expected since mucin is negatively charged due to the sulfuric acid and sulfate content and has exposed hydrophobic domains (1), it is anticipated that phage peptides that penetrate through mucin are hydrophilic and will have minimal positively charged residues (33). These properties are similar to net neutral, hydrophilic PEG-coated particles used for transmucosal delivery (13, 15, 21).

To confirm that identified peptides present on M13 have minimal mucin binding and achieve mucin penetration, we tested their transport through mucin compared with negative control wild-type M13. Here, clone TVRTSAD transports through mucin compared with wild-type, suggesting that the peptide improves transport. Except for the peptide sequence displayed on the N-terminus of p3 of TVRTSAD, both TVRTSAD and wild-type phage are genetically and phenotypically indistinguishable with identical coat proteins, which suggest that the peptide is primarily responsible for enhanced transport. The primary sequence of the peptide and the Kyte-Doolittle hydropathy plot suggest that the hydrophilic character of peptide may reduce phage adhesion to the hydrophobic domains of mucin and permit the phage to diffuse more rapidly. Using the method developed by Hu et al., we quantified the diffusive behavior of phage in mucin. TVRTSAD has an apparent diffusivity of 2.81*10^−9^ cm^2^/s, which is approximately over two-fold more than wild-type control. While these experiments were done in hyperconcentrated 8% mucin, the apparent diffusivities are the same order of magnitude as observed with diffusion of similarly structured filamentous fNEL phage through 1% agarose with and without *E. coli* (41, 43). fNEL exhibited apparent diffusivity of 2.6*10^−9^ cm^2^/s and 6.8*10^−8^ cm^2^/s through 1% and 0% agarose, respectively (41). The filamentous morphology decreased apparent diffusion of fNEL compared with icosahedral T4; It is hypothesized that the filamentous structure of fNEL more easily aggregates and gets trapped compared to smaller T4 (41). While selected phage demonstrates significant yet modest increase in diffusivity compared to negative control and is comparable to other filamentous phage diffusing through less concentrated agarose barriers, the diffusivity is less than observed with smaller icosahedral T4 or other PEG-coated particles (13, 41). While these effects to slow diffusivity are feasible, M13 is a flexible rod (48) able to take multiple conformations. M13 has a radius of gyration of ~140 nm (49), which is less than its length (~880 nm) yet greater than its diameter (~6-7 nm). The flexibility and ability to take multiple conformations potentially allows for peptide to reduce interaction with mucin to help facilitate transport.

Finally, we wanted to confirm that selected peptides were functional without display on the phage and could facilitate transport of small molecules as conjugates through the mucin barrier. FITC-labeled peptides were incubated in mucin and control saline to determine hindered and unhindered diffusion, respectively. All peptide conjugates and fluorescein salt exhibited decreased diffusion in mucin compared to saline. Fluorescein salt had 3-fold decreased diffusion, whereas peptide conjugates had approximately 4- to 5-fold reduced diffusivity. This suggests mucin, regardless or pore size, interacts and filters away small molecules through intermolecular interactions, as seen in previous studies with other small molecule drugs (50). Interestingly, the selected peptides demonstrated better effective diffusivity (D_PBS_/D_mucus_) than control cell-penetrating peptide hepta-arginine, which has hindered diffusion in mucus most likely due to charge interactions with net-negative charge mucin. This suggests that the amino acid composition can impact transport. Others have demonstrated that differences in amino acid sequence and composition can impact transport through mucin (16, 33). Li et al. demonstrated that net neutral charged peptides demonstrated improved transport compared to net positive and negative charged peptides; by altering the sequence of positive and negative charges could also impact the binding and penetration through the mucin barrier (33).

## CONCLUSIONS

We have developed a high-throughput screening assay using phage display libraries to identify peptides that reduce phage-mucin interactions and improve diffusivity of phage through hyperconcentrated mucin. Phage libraries provide a large collection of peptides with diverse chemistries (~10^9^ diversity) from which we can screen peptides that exhibit mucus-penetrating or muco-inert properties and may serve as a potential “coating” for drug delivery systems to improve transport through mucus barriers. From these initial screens, validated peptides were neutral to slightly negatively charged and are hydrophilic. Interestingly, these findings are consistent with previous work using low molecular weight, densely coated PEG nanoparticles as muco-inert particles for mucus penetration in multiple tissues (12, 13, 18, 20). While this study focused on high concentration of reconstituted mucin, future studies will require the use of more relevant concentrations and composition of mucus present in tissues and in diseases to truly recapitulate the (patho)physiology to understand and achieve mucus penetration (1, 15, 29, 30, 35). The selected phage clone exhibited improved diffusivity through high percentage mucin compared to control. To improve diffusivity, peptides could be engineered onto the p8 coat of M13, which has 2700 copies for more multivalent display and surface area to probe interactions with mucus. Alternatively, due to the “bulky” length of the filamentous M13 (~880 nm), further work to improve diffusivity could involve engineering peptides onto smaller viruses or functionalize onto polymeric 100 and 200 nm nanoparticles traditionally used in mucus penetrating delivery (13, 17–19). Out of the context of phage, the select peptides improved diffusivity compared to control positively charged peptides. However, they did not necessarily improve diffusion compared to fluorescein salt but possessed a higher molecular weight. Using next-generation sequencing, it may be possible to identify from a larger population of sequences additional peptides that may offer even further improvement on diffusivity (51, 52). While we identified a few peptides that did not share sequence homology with each other, with additional rounds of panning and sequence analysis we can begin to identify convergent peptide motifs that may be responsible for conferring optimal muco-inert properties. Through selective, systematic mutagenesis of amino acids of these peptides, it may be possible to dissect which amino acids (and their functionalities) are responsible for penetration through mucin. While we evaluated diffusivity of peptide conjugates to fluorophores, these muco-inert peptides could be conjugated to small molecule drugs previously impermeable to mucus penetration such as tobramycin and hydrocortisone or serve as potential alternative coatings for mucus penetrating drug delivery systems. From these insights, the promise of mucus-penetrating delivery can be realized.

## References

1. Cone RA. Barrier properties of mucus. Advanced drug delivery reviews. 2009;61(2):75–85.

2. Thornton DJ, Sheehan JK. From mucins to mucus: toward a more coherent understanding of this essential barrier. Proceedings of the American Thoracic Society. 2004;1(1):54–61.

3. Linden SK, Sutton P, Karlsson NG, Korolik V, McGuckin MA. Mucins in the mucosal barrier to infection. Mucosal immunology. 2008;1(3):183–97.

4. Niibuchi JJ, Aramaki Y, Tsuchiya S. Binding of Antibiotics to Rat Intestinal Mucin. Int J Pharm. 1986;30(2-3):181–7.

5. Hunt BE, Weber A, Berger A, Ramsey B, Smith AL. Macromolecular mechanisms of sputum inhibition of tobramycin activity. Antimicrobial agents and chemotherapy. 1995;39(1):34–9.

6. Komiya I, Park JY, Kamani A, Ho NFH, Higuchi WI. Quantitative Mechanistic Studies in Simultaneous Fluid-Flow and Intestinal-Absorption Using Steroids as Model Solutes. International Journal of Pharmaceutics. 1980;4(3):249–62.

7. Macadam A. The Effect of Gastrointestinal Mucus on Drug Absorption. Advanced drug delivery reviews. 1993;11(3):201–20.

8. Maher S, Mrsny RJ, Brayden DJ. Intestinal permeation enhancers for oral peptide delivery. Advanced drug delivery reviews. 2016;106(Pt B):277–319.

9. Zhang X, Wu W. Ligand-mediated active targeting for enhanced oral absorption. Drug Discov Today. 2014;19(7):898–904.

10. Serra L, Domenech J, Peppas NA. Engineering design and molecular dynamics of mucoadhesive drug delivery systems as targeting agents. European journal of pharmaceutics and biopharmaceutics: official journal of Arbeitsgemeinschaft fur Pharmazeutische Verfahrenstechnik eV. 2009;71(3):519–28.

11. des Rieux A, Pourcelle V, Cani PD, Marchand-Brynaert J, Preat V. Targeted nanoparticles with novel non-peptidic ligands for oral delivery. Advanced drug delivery reviews. 2013;65(6):833–44.

12. Dawson M, Wirtz D, Hanes J. Enhanced viscoelasticity of human cystic fibrotic sputum correlates with increasing microheterogeneity in particle transport. The Journal of biological chemistry. 2003;278(50):50393–401.

13. Lai SK, O’Hanlon DE, Harrold S, Man ST, Wang YY, Cone R, et al. Rapid transport of large polymeric nanoparticles in fresh undiluted human mucus. Proceedings of the National Academy of Sciences of the United States of America. 2007;104(5):1482–7.

14. Cu Y, Saltzman WM. Controlled surface modification with poly(ethylene)glycol enhances diffusion of PLGA nanoparticles in human cervical mucus. Molecular pharmaceutics. 2009;6(1):173–81.

15. Crater JS, Carrier RL. Barrier properties of gastrointestinal mucus to nanoparticle transport. Macromolecular bioscience. 2010;10(12):1473–83.

16. Lieleg O, Vladescu I, Ribbeck K. Characterization of particle translocation through mucin hydrogels. Biophysical journal. 2010;98(9):1782–9.

17. Schuster BS, Suk JS, Woodworth GF, Hanes J. Nanoparticle diffusion in respiratory mucus from humans without lung disease. Biomaterials. 2013;34(13):3439–46.

18. uk JS, Lai SK, Wang YY, Ensign LM, Zeitlin PL, Boyle MP, et al. The penetration of fresh undiluted sputum expectorated by cystic fibrosis patients by non-adhesive polymer nanoparticles. Biomaterials. 2009;30(13):2591–7.

19. Tang BC, Dawson M, Lai SK, Wang YY, Suk JS, Yang M, et al. Biodegradable polymer nanoparticles that rapidly penetrate the human mucus barrier. Proceedings of the National Academy of Sciences of the United States of America. 2009;106(46):19268–73.

20. Wang YY, Lai SK, Suk JS, Pace A, Cone R, Hanes J. Addressing the PEG mucoadhesivity paradox to engineer nanoparticles that “slip” through the human mucus barrier. Angew Chem Int Ed Engl. 2008;47(50):9726–9.

21. Lautenschlager C, Schmidt C, Lehr CM, Fischer D, Stallmach A. PEG-functionalized microparticles selectively target inflamed mucosa in inflammatory bowel disease. European journal of pharmaceutics and biopharmaceutics: official journal of Arbeitsgemeinschaft fur Pharmazeutische Verfahrenstechnik eV. 2013;85(3 Pt A):578–86.

22. Dams ET, Laverman P, Oyen WJ, Storm G, Scherphof GL, van Der Meer JW, et al. Accelerated blood clearance and altered biodistribution of repeated injections of sterically stabilized liposomes. The Journal of pharmacology and experimental therapeutics. 2000;292(3):1071–9.

23. Abu Lila AS, Kiwada H, Ishida T. The accelerated blood clearance (ABC) phenomenon: clinical challenge and approaches to manage. Journal of controlled release: official journal of the Controlled Release Society. 2013;172(1):38–47.

24. Richter AW, Akerblom E. Antibodies against polyethylene glycol produced in animals by immunization with monomethoxy polyethylene glycol modified proteins. International archives of allergy and applied immunology. 1983;70(2):124–31.

25. Cheng TL, Wu PY, Wu MF, Chern JW, Roffler SR. Accelerated clearance of polyethylene glycol-modified proteins by anti-polyethylene glycol IgM. Bioconjugate chemistry. 1999;10(3):520–8.

26. Mishra S, Webster P, Davis ME. PEGylation significantly affects cellular uptake and intracellular trafficking of non-viral gene delivery particles. European journal of cell biology. 2004;83(3):97–111.

27. Remaut K, Lucas B, Braeckmans K, Demeester J, De Smedt SC. Pegylation of liposomes favours the endosomal degradation of the delivered phosphodiester oligonucleotides. Journal of controlled release: official journal of the Controlled Release Society. 2007;117(2):256–66.

28. Hatakeyama H, Akita H, Kogure K, Oishi M, Nagasaki Y, Kihira Y, et al. Development of a novel systemic gene delivery system for cancer therapy with a tumor-specific cleavable PEG-lipid. Gene therapy. 2007;14(1):68–77.

29. Murgia X, Loretz B, Hartwig O, Hittinger M, Lehr CM. The role of mucus on drug transport and its potential to affect therapeutic outcomes. Advanced drug delivery reviews. 2018;124:82–97.

30. Leal J, Smyth HDC, Ghosh D. Physicochemical properties of mucus and their impact on transmucosal drug delivery. Int J Pharm. 2017;532(1):555–72.

31. Olmsted SS, Padgett JL, Yudin AI, Whaley KJ, Moench TR, Cone RA. Diffusion of macromolecules and virus-like particles in human cervical mucus. Biophysical journal. 2001;81(4):1930–7.

32. Wada A, Nakamura H. Nature of the charge distribution in proteins. Nature. 1981;293(5835):757–8.

33. Li LD, Crouzier T, Sarkar A, Dunphy L, Han J, Ribbeck K. Spatial configuration and composition of charge modulates transport into a mucin hydrogel barrier. Biophysical journal. 2013;105(6):1357–65.

34. Teesalu T, Sugahara KN, Ruoslahti E. Mapping of vascular ZIP codes by phage display. Methods Enzymol. 2012;503:35–56.

35. Lai SK, Wang YY, Wirtz D, Hanes J. Micro- and macrorheology of mucus. Advanced drug delivery reviews. 2009;61(2):86–100.

36. Green MR, Sambrook J, Sambrook J. Molecular cloning: a laboratory manual. 4th ed. Cold Spring Harbor, N.Y.: Cold Spring Harbor Laboratory Press; 2012.

37. Henderson AG, Ehre C, Button B, Abdullah LH, Cai LH, Leigh MW, et al. Cystic fibrosis airway secretions exhibit mucin hyperconcentration and increased osmotic pressure. The Journal of clinical investigation. 2014;124(7):3047–60.

38. Matsui H, Wagner VE, Hill DB, Schwab UE, Rogers TD, Button B, et al. A physical linkage between cystic fibrosis airway surface dehydration and Pseudomonas aeruginosa biofilms. Proceedings of the National Academy of Sciences of the United States of America. 2006;103(48):18131–6.

39. McGill SL, Smyth HD. Disruption of the mucus barrier by topically applied exogenous particles. Molecular pharmaceutics. 2010;7(6):2280–8.

40. Kyte J, Doolittle RF. A simple method for displaying the hydropathic character of a protein. J Mol Biol. 1982;157(1):105–32.

41. Hu J, Miyanaga K, Tanji Y. Diffusion properties of bacteriophages through agarose gel membrane. Biotechnology progress. 2010;26(5):1213–21.

42. Ortiz ME, Endy D. Engineered cell-cell communication via DNA messaging. Journal of biological engineering. 2012;6(1):16.

43. Hu J, Miyanaga K, Tanji Y. Diffusion of bacteriophages through artificial biofilm models. Biotechnology progress. 2012;28(2):319–26.

44. Hopp TP, Woods KR. Prediction of protein antigenic determinants from amino acid sequences. Proceedings of the National Academy of Sciences of the United States of America. 1981;78(6):3824–8.

45. Desai M, Vadgama P. Estimation of effective diffusion coefficients of model solutes through gastric mucus: assessment of a diffusion chamber technique based on spectrophotometric analysis. Analyst. 1991;116(11):1113–6.

46. Sidhu SS. Phage display in biotechnology and drug discovery. Boca Raton: CRC Press/Taylor & Francis; 2005. xviii, 748 p. p.

47. Matochko WL, Cory Li S, Tang SK, Derda R. Prospective identification of parasitic sequences in phage display screens. Nucleic Acids Res. 2014;42(3):1784–98.

48. Khalil AS, Ferrer JM, Brau RR, Kottmann ST, Noren CJ, Lang MJ, et al. Single M13 bacteriophage tethering and stretching. Proceedings of the National Academy of Sciences of the United States of America. 2007;104(12):4892–7.

49. Ghosh D, Lee Y, Thomas S, Kohli AG, Yun DS, Belcher AM, et al. M13-templated magnetic nanoparticles for targeted in vivo imaging of prostate cancer. Nat Nanotechnol. 2012;7(10):677–82.

50. Bhat PG, Flanagan DR, Donovan MD. Drug diffusion through cystic fibrotic mucus: steady-state permeation, rheologic properties, and glycoprotein morphology. J Pharm Sci. 1996;85(6):624–30.

51. Derda R, Tang SK, Li SC, Ng S, Matochko W, Jafari MR. Diversity of phage-displayed libraries of peptides during panning and amplification. Molecules. 2011;16(2):1776–803.

52. Liu GW, Livesay BR, Kacherovsky NA, Cieslewicz M, Lutz E, Waalkes A, et al. Efficient Identification of Murine M2 Macrophage Peptide Targeting Ligands by Phage Display and Next-Generation Sequencing. Bioconjugate chemistry. 2015;26(8):1811–7.

